# Performance of eDNA capture methods for monitoring fish biodiversity in a hyper-tidal estuary

**DOI:** 10.1101/2024.12.20.629404

**Authors:** Jake M. Jackman, Naiara Guimarães Sales, Chiara Benvenuto, Andrea Drewitt, Andrew Wolfenden, Peter E. Robins, Ilaria Coscia, Allan D. McDevitt

## Abstract

Environmental DNA (eDNA) has become an established and efficient method for monitoring biodiversity in aquatic systems. However, there is a need to compare and standardise sampling methods across ecosystem types, particularly complex ecosystems such as estuaries where unique challenges for monitoring fish populations are present due to fluctuating environmental factors. Here, we compare fish biodiversity metrics obtained from eDNA metabarcoding data using four different eDNA filtering methods: three manual filtering methods with different pore sizes (0.45, 1.2 and 5 µm) and a newly established passive method, the metaprobe. The study was applied across a salinity gradient in a hyper-tidal estuarine ecosystem. Overall, 44 fish species were detected across the four methods used. The 0.45 µm filter recovered the highest richness (39 species), then the metaprobe method (35), followed by the 1.2 µm (34) and 5 µm (33) filters. Filter performance between salinity gradients revealed that the 0.45 µm and the 1.2 µm methods recovered the highest species richness across all sampled zones. The 0.45 µm also had the most consistent detection probabilities using representative species from each zone. While the 0.45 µm method appeared to be the optimal method, each of the methods can be considered as a viable and comparable option for biomonitoring in dynamic ecosystems such as estuaries and rivers. In particular, the passive metaprobe (used in a freshwater system for the first time here) performed well in comparison to the manual filtering methods despite a short deployment time. This study provides critical insights for optimising fish biodiversity assessments using eDNA metabarcoding in estuarine ecosystems, providing a valuable framework for future monitoring efforts in similar systems worldwide.

## Introduction

Monitoring organisms in complex environments, such as fishes in estuarine systems, poses challenges associated with different sampling methodologies (Alenzi, 2024). Traditional fish monitoring relies on the capture, visual identification and counting of specimens (Radinger et al., 2019; Franco et al., 2021), using approaches such as gill/seine-netting, electrofishing, beam trawling and recreational angling (Magaju et al., 2023; Baldino et al., 2018). However, these methodologies can be invasive and expensive to undertake. The application of DNA-based approaches, particularly environmental DNA (eDNA) metabarcoding - the simultaneous amplification/sequencing of DNA from multiple species - has become an established and efficient method for monitoring biodiversity (Taberlet et al., 2012; Ogden, 2022). The value that eDNA metabarcoding can provide as a monitoring tool has been well evidenced in a range of aquatic ecosystems worldwide (Rees et al., 2014; McDevitt et al., 2019; Valentin et al., 2020; Sales et al., 2021), including analyses of seasonal dynamics of fish communities in freshwater river systems (Milhau et al., 2021), investigations of spatio-temporal variability of eDNA in freshwater lakes (Herve et al., 2022), description of fish communities in marine biodiversity hotspots (Valdivia-Carillo et al., 2021), and the evaluation of fish biodiversity in estuaries (Zainal Abindin et al., 2022).

While eDNA analysis has been shown to be a powerful tool for biodiversity monitoring, it is not without its challenges, particularly when sampling in aquatic systems characterised by complex mixing dynamics and density gradients, such as estuaries (Sanches & Schreier, 2020; DiBattista et al., 2022). In contrast to oceans, lakes and rivers, where the water chemistry tends to be more homogeneous, estuaries encompass both marine and freshwater - resulting in strong and variable horizontal salinity gradients, with the added complexity of resuspended sediments, which can cause the filters to clog (Williams et al., 2017). The most common tools used to capture eDNA are filter membranes enclosed within a plastic cartridge, attached to a manual or mechanical syringe/pump which forces water through the membrane (Deiner et al., 2015; Miya et al., 2016; Capo et al., 2020). These filters come in various pore sizes, i.e., the size of microscopic holes across the surface of the membrane, measured in micrometres (μm). The diameter of these pores allows water to pass through while capturing genetic material on the surface of the membrane. The pore sizes commonly selected for eDNA studies range from 0.22 μm to 5 μm (Turner et al., 2014; Thomas et al., 2018). Larger pores are expected to filter more water before becoming clogged and thus yield a higher concentration of DNA (Kumar et al., 2022). However, this can introduce challenges in terms of capturing low-concentration DNA, as larger-pore filters may allow smaller DNA fragments to pass through, reducing detection sensitivity (Jo, Takao & Minamoto, 2022).

In highly dynamic and turbid environments such as tidally energetic estuaries, the suspended particles within a water sample can quickly obstruct the filter and inhibit the filtration process regardless of pore size (Barnes et al., 2021; Hallam et al., 2021). To circumvent the challenges associated with manual filtration, the use of passive samplers has been proposed (Bessey et al., 2021): absorbent materials are placed within the water source and passively accumulate eDNA particles over varied time frames (Jeunen et al., 2022; Verdier et al., 2022; Chen et al., 2024). Recently, a new passive sampler, the metaprobe (i.e., a 3D-printed spherical probe in which absorbent gauze is held), has been tested (Maiello et al., 2022). However, its efficiency has not been evaluated in estuarine or freshwater environments, and so far, has exclusively been deployed within marine environments, typically placed inside nets, and cast out from fishing vessels (Maiello et al., 2024; Sbrana et al., 2024). Currently, there is no specific or standardised method for filtering and capturing eDNA (Hirsch et al., 2024), which presents an ongoing challenge to the application of eDNA as a standardised monitoring method. This lack of standardisation leads to a wide diversity of approaches, with new studies potentially implementing protocols that may unknowingly be sub-optimal for their specific target systems (Sanches & Schreier, 2020). Such variability can compromise the quality of results, particularly in environments with unique challenges, like estuaries characterised by high turbidity and dynamic water chemistry. This issue reinforces the need to develop and evaluate best practices tailored to different environmental conditions. Establishing standardised protocols would not only enhance the reliability and quality of eDNA studies but also accelerate the adoption of this technique as a robust tool for biodiversity monitoring in complex ecosystems.

The Mersey estuary (UK) is a complex aquatic ecosystem characterised by strong, dynamic tidal behaviour (Bowden, 1963) and was once home to thriving populations of Atlantic salmon (*Salmo salar*), eel (*Anguilla anguilla*), smelt (*Osmerus eperlanus*), lampreys (*Lampetra fluviatilis* and *Petromyzon marinus*) and brown trout (*Salmo trutta*), all of which are present on the UK Biodiversity Action Plan (BAP) priority fish species list (Joint Nature Conservation Committee, 2007). This system suffered severe water quality degradation during the Industrial Revolution, earning the reputation of one of the most polluted rivers in Europe (Jones, 2006). During this time, most fish populations within the system collapsed, and what was once a thriving fisheries ground became biologically barren (Jones, 2000). However, the implementation of new water quality legislation in the 1980s, through the efforts of the Mersey Basin Campaign, an initiative implemented to promote an ecological recovery of the Mersey (Kim & Batey, 2021), led to significant improvements in the habitat quality, followed by sightings of Atlantic salmon within the estuary in 1999 after decades of absence (Jones, 2006; Buysse et al., 2008; Ikedaishi et al., 2012). In the case of salmon, given the low estimated numbers of fish that return to spawn as part of their life cycle (Mawle & Milner, 2003), relying on catching physical specimens with traditional monitoring might not be the most effective method.

In this study, eDNA metabarcoding was applied as a biomonitoring tool in the Mersey estuary: four different eDNA capturing methods were tested across ten sites which encompass the lower, central and upper zones of the estuary, with each zone featuring differing water chemistry dynamics (marine, brackish and freshwater). The aim was to compare the overall performance of each capture method in terms of multiple fish biodiversity diversity metrics across different physical habitats and to determine which one could be routinely used for future biodiversity monitoring campaigns of highly heterogeneous estuaries. Overall, our goal is to inform future biomonitoring strategies for tidal estuaries, where tides, sediment resuspension and variable salinity gradients shape biodiversity distributions. The findings could inform monitoring plans in similar habitats worldwide and highlight which factors should be considered when designing eDNA biomonitoring campaigns.

## Methods and materials

### Study area

The Mersey estuary (UK), extends ∼60 km from the upper tidal limit near Warrington to Liverpool bay in the Irish Sea (Fig. 1). The estuary is hyper-tidal with a spring tidal range of ∼10 m at the mouth (∼4 m during neap tides), and is semi-diurnal with two similar tidal cycles per day (Lane, 2004). Saltwater from the Irish Sea mixes with freshwater inputs, mainly from the river Mersey, where the annual mean flow is approximately ∼37 m³/s. Low flows, represented by the 95th percentile exceedance (flows occurring 95% of the time) value, are around ∼ 9.4 m³/s, while high-flow conditions, corresponding to the 5th percentile exceedance (flows occurring 5% of the time), reach approximately ∼93 m³/s (National River Flow Archive, 2024). These factors contribute to a highly energetic and well-mixed estuary that shapes species distributions throughout the river-to-coast continuum (Elliot & McLusky, 2002).

**Figure 1.**
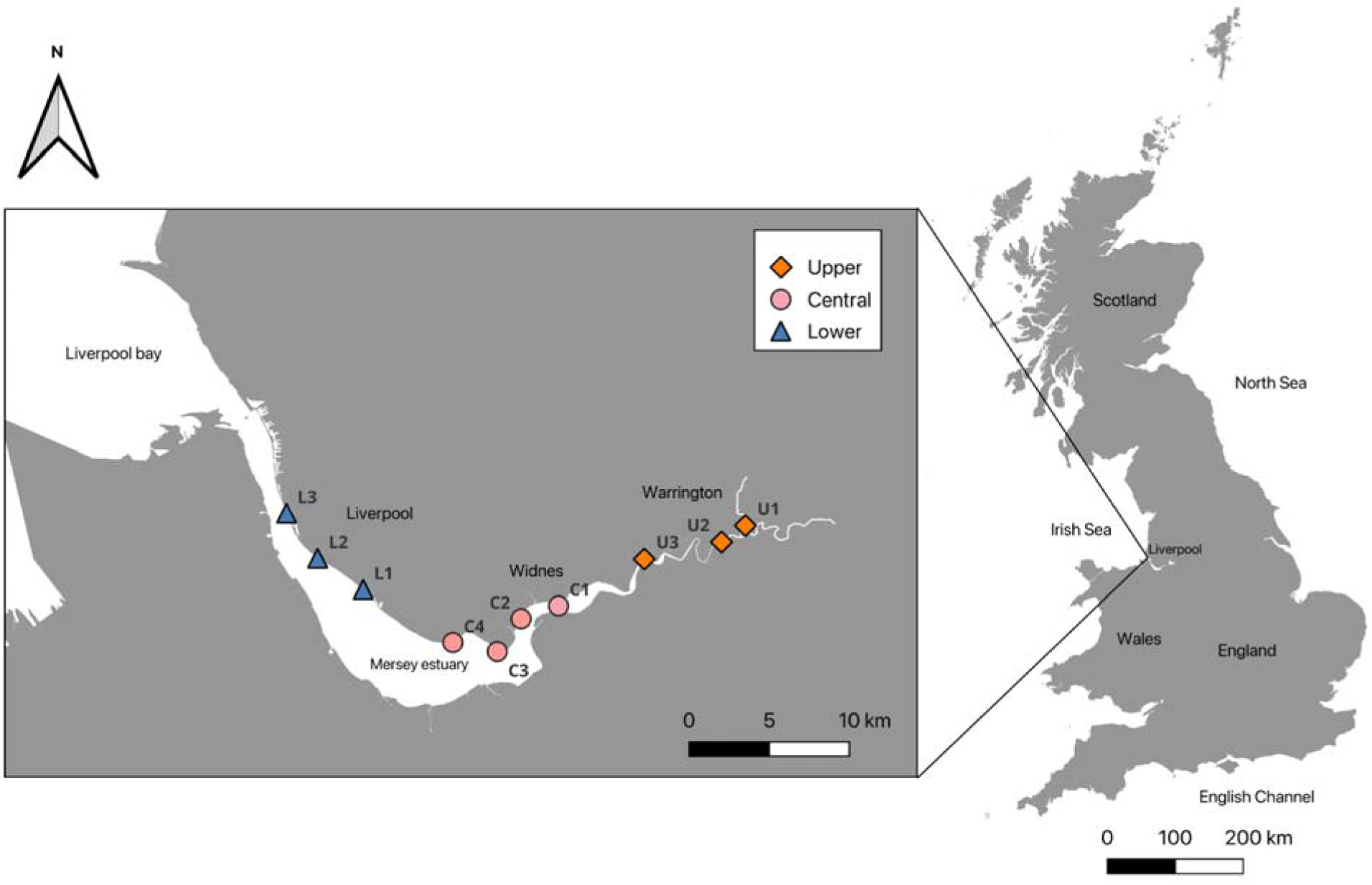
Sampling sites on the Mersey estuary: three in the upper zone (orange diamonds), four in the central zone (pink circles) and three in the lower zone (blue triangles).

For this study, 10 sampling locations on the northern banks of the estuary were selected, spanning the three distinct zones within the system. In the upper limits of the estuary, which are predominately freshwater locations, three sites were sampled (U1, U2 and U3). The central areas of the estuary form an Estuarine Turbidity Maximum (ETM), a zone of highest turbidity resulting from turbulent resuspension of sediment (Geyer, 1993). This zone creates a large mixing section as the marine water converges with the freshwater. In total, four sample sites were selected here (C1, C2, C3 and C4) as this area is more prominent (∼5 km in width) and has added complexity resulting from the ETM. The lower estuary, which is primarily a marine zone, had another three sample sites designated (L1, L2 and L3) (Fig. 1).

### eDNA capture methods

Four distinct methods were evaluated including syringe filters with pore sizes of 0.45 μm (Sterivex™), 1.2 μm (Whatman™), 5 μm (KX Nylon©), and a passive method, the metaprobe (Fig. 2).

**Figure 2.**
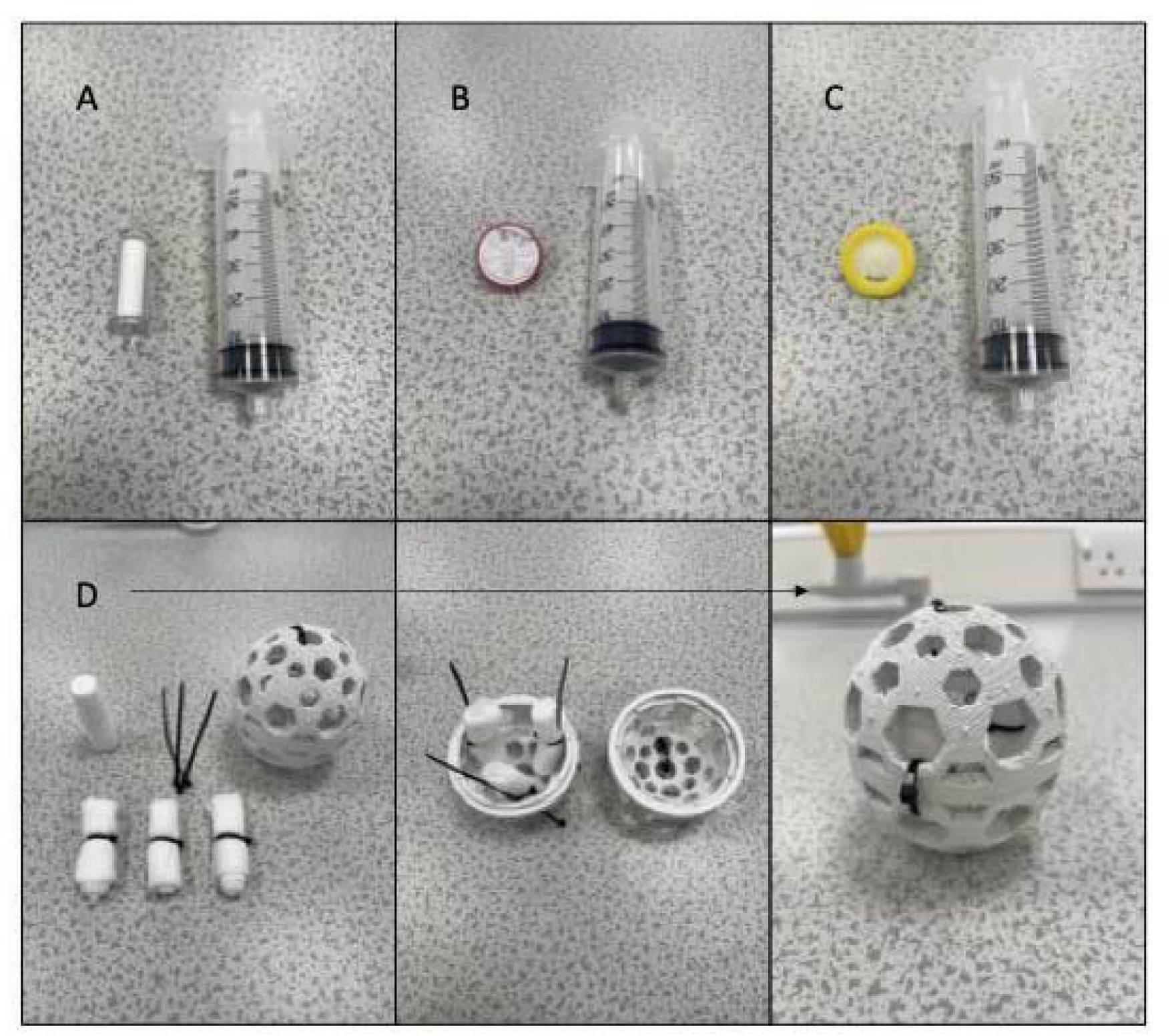
The four different sampling methods tested in the study, including Sterivex 0.45 µm (A), Whatman 1.2 µm (B), KX Nylon 5 µm (C) and the metaprobe (D).

Samples were collected during November and December 2022 and January 2023, during three sampling days per month, each time during high tide due to multiple sampling locations being frequently dry at low tides. During each month, three replicates for each method were taken at each of the 10 sites, for a total of 360 eDNA samples collected across the experiment. For the three methods using syringe filters, water samples were collected in individual 1 L sterilised buckets, using one bucket per replicate. A single 1 L field blank water replicate was taken at the beginning of each sampling day using sealed bottled water. Each of the syringe-based filters used a 60 ml syringe to manually pass the water through the filter membrane. Filtration of the water samples was performed on-site immediately following collection. Once each syringe filter replicate became clogged, volume filtered was recorded, and the filter was sealed in an individual sterile air-tight bag and stored at −20°C in the laboratory until processed. The metaprobe method was prepared in a dedicated eDNA clean room at the University of Salford before fieldwork. Individual rolls (three rolls representing three replicates) of identical size (10 × 10 cm) medical grade gauze were attached to the inside walls of the metaprobe sphere, with sterilised cable ties. For each metaprobe, three 50 ml falcon tubes containing 99% ethanol were prepared for storage of the gauze after sampling. The prepared metaprobe and falcon tubes were sealed in air-tight sterile bags until deployment at their respective sampling locations, immediately after completion of the syringe filtering method. The metaprobe was attached to a 20 m rope using a carabiner (Fig. S1) and cast out into the water for five minutes at each site. Prior to the metaprobe deployment, a single field blank gauze replicate was placed in a falcon tube containing 99% ethanol at the beginning of each sampling day. Immediately following deployment, the metaprobe was carefully cut open and each of the rolls of gauze was placed in a pre-prepared individual falcon tube and stored at −20°C until DNA extraction. The five-minute duration was determined based on the average time to filter one replicate for each of the other methods, therefore creating an equivalent sampling duration across all methods tested.

### DNA extraction and amplification

All DNA extractions were performed in a dedicated eDNA clean room, and decontaminated via UV for at least six hours before extractions by personnel wearing full PPE. DNA extractions followed the Mu-DNA protocol (Sellers et al., 2018) tailored for water samples (for the syringe-based filtering methods) and soil samples (for the metaprobe capturing method). The choice of the soil extraction method was made to account for the unique characteristics of the metaprobe, as they collect and encapsulate particulates, commonly suspended sediments, creating a sample composition more akin to soil. In total 360 eDNA samples and 36 negative controls, including field and laboratory blanks, were processed.

Two different primer sets were tested: MiFish-U/E and Tele02. The MiFish-U/E primers target a hypervariable region of the 12S rRNA gene (163–185 bp) and have been used extensively in freshwater settings (Miya et al., 2015); the Tele02 primers amplify a ∼167 bp fragment of the 12S gene, specifically targeting teleost fishes (Taberlet et al., 2018). Although both primers successfully amplified the target gene regions, the MiFish primers, when used on brackish and marine water samples (i.e., samples collected in the central and lower zone), amplified a predominant non-target fragment of unknown origin (Fig. S2) whereas the Tele02 primers mainly amplified the fish-specific target fragment (Fig. S2). Due to the lack of consistency with the MiFish primers across all sampled zones, extracted DNA was amplified using the fish-specific Tele02 primers (Taberlet et al., 2018). Both a positive (*Hoplias malabaricus,* a neotropical freshwater species absent in the UK) and a negative PCR control (i.e., PCR reaction using nuclease-free water instead of eDNA) were included to account for possible tag jumping and contamination.

PCRs were prepared across six equally balanced libraries (90 rolls of gauze – 15 per library, 270 syringe filter samples – 45 per library, 12 field blanks – 2 per library, 12 extraction blanks – 2 per library, 12 PCR negative controls – 2 per library and 12 PCR positive controls – 2 per library; 68 samples total per library), and PCR was performed in triplicate for each library. Each sample was amplified using the Teleo02 primers including a unique 8 bp oligo-tag attached to the forward and reverse primers and a variable number (2-4) of leading Ns (fully degenerate positions) to increase variability in amplicon sequences. PCR amplification was conducted using a single-step protocol to minimise bias in individual reactions. The PCR reaction consisted of a total volume of 20 µL including 10 µl AmpliTaq Gold™ 360 Master Mix (1X; Applied Biosystems); 0.16 µl of BSA (5 mg); 1 µl of each of the two primers (5 µM); 5.84 µl of ultra-pure water and 2 µl of eDNA template. All PCR amplification of libraries was performed under the following thermocycling conditions: 95°C for 10 min, followed by 40 cycles of 95°C for 30 s, 60°C for 45 s, and 72°C for 30 s, and a final elongation of 72°C for 5 min.

Replicates were then pooled, and samples were visualised on a 1.2% agarose gel stained with GelRed® to check for successful amplification of target fragments. PCR products were then purified with HighPrep™ PCR Clean-up System magnetic beads using a 1.1× ratio for a left-sided size selection. The purified libraries were visualised on the Agilent 2200 TapeStation using High Sensitivity D1000 ScreenTape (Agilent Technologies). This indicated secondary non-target products on the right side of the target fragment which was removed by right-sided size selection (0.8x ratio for all libraries).

Size selected DNA was quantified using a Qubit™ 4.0 fluorometer with the Qubit™ dsDNA HS Assay Kit (Invitrogen). Based on the total DNA concentration, each library was diluted to 20 ng/ul at a volume of 50 µl for library preparation. End repair, adapter ligation and library PCR amplification were performed using the KAPA HyperPrep Kit according to the manufacturer’s protocol. Libraries were quantified using quantitative PCR (qPCR) on a MIC qPCR system (Bio Molecular Systems) with the NEBNext® Library Quant Kit for Illumina® (New England Biolabs). Libraries 1 & 2, 3 & 4 and 5 & 6 were pooled in equimolar concentrations to create a final three pools. All libraries were sequenced at 9 pM (3 sequencing runs total) on Illumina® MiSeq™ Reagent v2 (300-cycle) kits (Illumina Inc.).

### Bioinformatic analysis

The bioinformatics analysis was carried out using OBITools 1.2.11 (Boyer et al., 2016). Read quality was checked with *fastqc* (Andrews, 2010) and low-quality ends were trimmed for downstream analysis. We used *illuminapairedend* to merge all paired reads with a quality score >30, and *ngsfilter* to demultiplex samples based on their unique barcodes. Sequences were filtered via *obigrep* to remove singletons and reads out of the expected length range (129–209 bp), and dereplicated via *obiuniq*. We removed chimeras with *uchime* (Edgar et al., 2011) and clustered the remaining sequences into Molecular Operational Taxonomic Units (MOTU) with *swarm* (Mahé et al., 2015) setting the threshold to d = 1. Sequences were assigned taxonomy information using a DNA reference library dataset for fish species of the UK, derived from the NCBI GenBank and Barcode of Life BOLD databases (Meta-Fish-Lib v255). The reference dataset includes species for both freshwater and marine UK species (Collins et al., 2021).

The final dataset was manually curated to improve taxonomic assignments by checking MOTUs identified at the genus level. This was due to a small number of species not being distinguished correctly, likely due to limitations within the reference database. For this reason, MOTUs assigned only at the genus level were checked against the NCBI nucleotide database for the confirmation of taxonomic assignment (Altschul et al., 1990). Poorly annotated MOTUs were defined as those with ambiguous assignments, such as sequences that were only classified to high taxonomic ranks (e.g., family or order) and other sequences which had been assigned to incorrect species. This revealed the correct identities of seven previously misassigned species or species in which the ID thresholds were unusually low, given that the species assigned are known to inhabit the study area. These were: the common roach *Rutilus rutilus*, dace *Leuciscus leuciscus* and common carp *Carpio carpio* in the Cyprinid order; the European flounder *Platichthys flesus,* common dab *Limanda limanda* and plaice *Pleuronectes platessa* in the Pleuronectiformes; and the whiting *Merlangius merlangus* in the Gadiformes.

Stringent filtering steps, performed in R v4.3.1 (R core team 2023), were applied to the final dataset to remove MOTUs/reads originating from sequencing errors or contamination to avoid false positives for the library using the ‘tidyverse’ (Wickham et al., 2019) and ‘dyplr’ (Wickham et al., 2023) packages. All non-fish reads were removed from the dataset, including non-target species (e.g., human and domestic species reads) and MOTUs that were likely to have been carried over from contamination. To remove contaminants, the maximum number of reads recorded in the controls (field collection blanks, DNA extraction blanks and PCR blanks) were removed from all samples. To address the potential of tag jumping, the sample data was examined to assess whether any positive control-specific MOTUs were present in any other samples, and all reads associated with the positive control were subtracted from the sample data. Finally, all MOTUs with < 5 reads were removed from the final dataset. The final taxonomic assignment was conducted according to current fixed general thresholds: MOTUs were assigned at the species level when matching the reference sequence with > 98% in line with other UK fish studies (Hallam et al., 2021; Rourke et al., 2022).

### Statistical analysis

All downstream analyses were performed in R v4.3.1 (R core team, 2023). The presence/absence data of species detected in each of the three sampling months were combined to create an overall dataset of species richness obtained from the total duration of the study sampling campaign.

To determine the overall richness obtained by each eDNA capturing method and the number of species detections shared, and exclusively detected by each sampling method, a box plot and Venn diagram were created using the R-packages: ‘ggplot2’ and ‘VennDiagram’. Additionally, a permutational multivariate analysis of variance (PERMANOVA) was performed to test for significant differences in species richness between the four sampling methods, followed by post-hoc pairwise PERMANOVA comparisons. Rarefaction and extrapolation curves with 95% confidence intervals were used to compare the overall number of species richness recovered by each method and sampling effort for the 10 sites sampled (R-package: ‘iNEXT’ v2.0.20; Chao et al., 2014).

The data was visualised using a combined grouped boxplot and violin plot showing overall richness by capture method within the three zones present within the study system (lower, central and upper). Statistical comparisons were then performed to distinguish meaningful increases or decreases in species richness between the four methods and across the three zones with a Generalised Linear Mixed Model (GLMM) using the function *glmer* from the R-package ‘lme4’ (v1.1-35; Bates et al., 2015). The GLMM was fitted using Maximum Likelihood Estimation (MLE) with Laplace Approximation (Raudenbush et al., 2000). The response variable, “Species Richness”, was modelled using a Poisson distribution with a logarithmic link function. “Site” was included as a random effect in the model to estimate the variability in Species Richness among different sites, beyond what can be explained by the nested fixed effects (Zone/Method). The model was formulated as follows: *Species Richness* ∼ *Zone/Method + (1*∣*Site).* The GLMM was implemented using a nested approach, to determine if species richness significantly increases or decreases when using a specific method inside and among the different zones (Method: filtering, designated by pore size of membrane: 0.45 µm, 1.2 µm, 5 μm or passive metaprobe). The GLMM requires a baseline for comparisons to be made (intercept), here, the intercept was set as the species richness recovered in each zone using the 0.45 µm pore size, therefore, all comparisons are made against the 0.45 µm method (e.g., the values associated with ‘Central: Method 1.2 µm’ are compared to ‘Central: Method 0.45 µm’).

To account for richness differences which may be due to non-standardised volumes of water filtered by each method, a Pearson’s correlation test between the volume of water filtered and species richness was performed. Scaling Ranked Subsampling (SRS; Beule & Karlovsky, 2020) was used to correct the species richness based on a normalised volume of water filtered across all samples. The normalised value (*Cmin*) was selected to be the lowest volume of water filtered between the 0.45 µm, 1.2 µm and 5 µm syringe filtration methods (*Cmin* = 180 ml). The metaprobe was excluded from this test as the water that has been processed by this method could not be quantified.

To demonstrate the relative efficacy of the four eDNA sampling methods, we performed species detection probability and species-specific cumulative detection probability analyses as used by Sales et al. (2020). Here, we performed these analyses on the four eDNA sampling methods, assessing the three replicates collected using three representative species of fish found within the eDNA dataset: European eel *Anguilla anguilla*, European sprat *Sprattus sprattus* and the common bream *Abramis brama*. These species were selected as representatives to provide us with coverage of species known to inhabit all three zonal gradients (European eel), only the lower zone (marine; European sprat) and only the upper zone (freshwater; common bream).

## Results

The resultant dataset obtained after quality-checking and filtering consisted of 6,114,309 reads in total (Table S1), which allowed the detection of 44 unique fish species (Table S2). We recovered 36 species (2,924,358 reads) in November 2022, 38 species (2,333,900 reads) in December 2022, and 33 species (856,046 reads) in January 2023.

The overall species richness detected for all months combined varied significantly across the sampling methods (Fig. 3A; PERMANOVA: *F* = 4.51, *p* = 0.005) with the highest average richness obtained by the 0.45 μm and the 1.2 μm filter methods (PERMANOVA: *F* = 0.1073, *p* = 0.879) and the lowest average richness obtained by the 5 μm filter method and the metaprobe (PERMANOVA: *F* = 0.281, *p* = 0.735).

**Figure 3.**
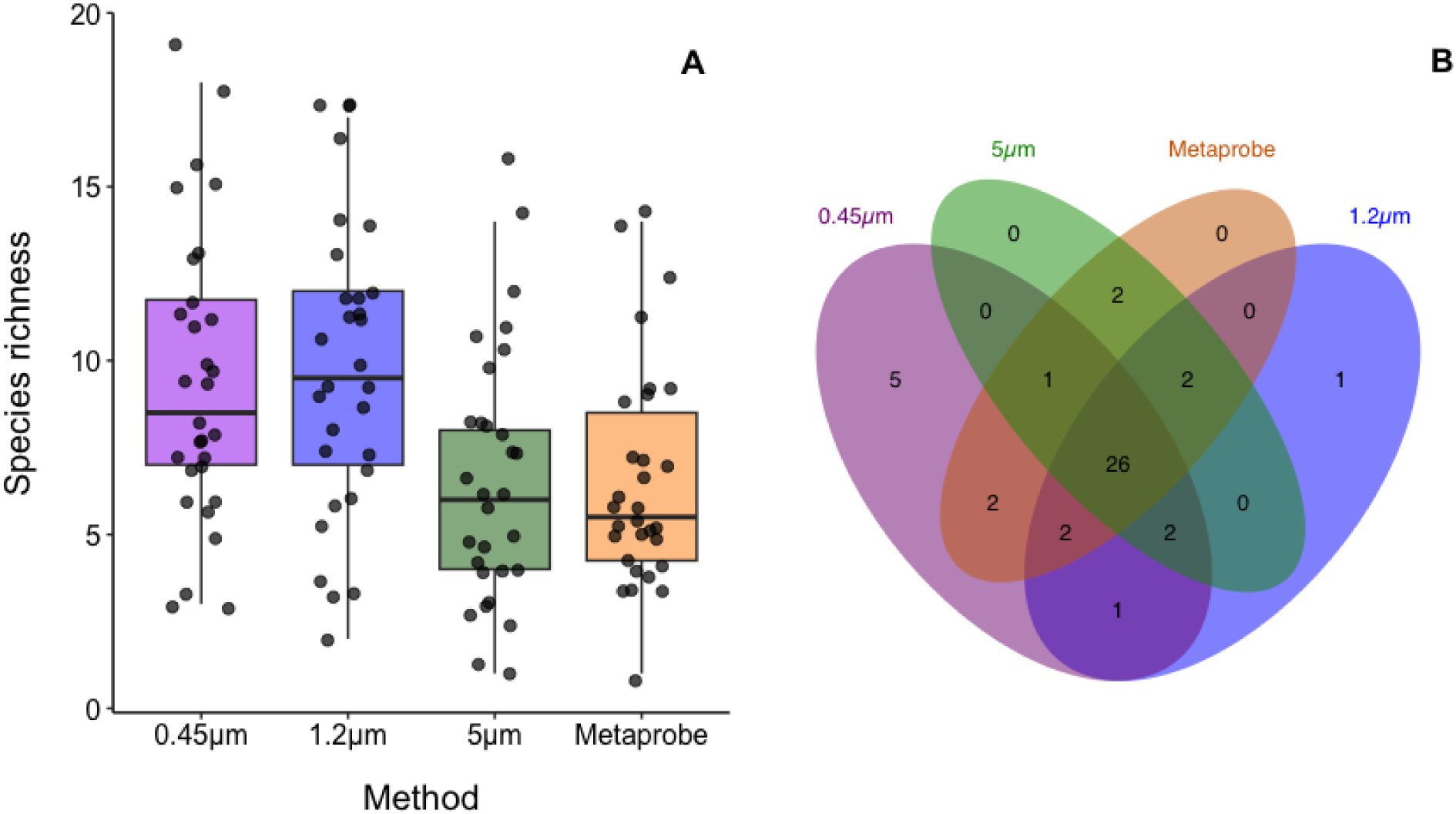
Overall species richness detected by each eDNA capturing method (designated by filtering pore size for the filtration methods; A) and the total, shared and exclusively detected numbers of species by each method (B).

The total number of species detected by each capture method can be observed in the Venn diagram (Fig. 3B), which also highlights the number of species exclusively detected for each method. Overall, 26 out of 44 (∼60%) species were detected by all four methods. The 0.45 μm filter detected 39 species (87% of the total), the 1.2 μm detected 34 species (75% of the total), the 5 μm detected 33 species (73% of the total) and the metaprobe detected 35 species (78% of the total). Of the 44, five (11%) were exclusively detected by the 0.45 μm, one (2%) exclusively by the 1.2 μm and neither the 5 μm nor the metaprobe exclusively detected any species (see Table S2 for details on which species were detected by each method).

Rarefaction and extrapolation curves show a higher projected asymptotic species richness for the 0.45 µm method, followed by the metaprobe, 1.2 µm, and 5 µm methods (Fig. 4). The extrapolation curves for the 1.2 µm, 5 µm and the metaprobe reach a plateau between ∼20-30 sites and the 0.45 µm at ∼40 sites.

**Figure 4.**
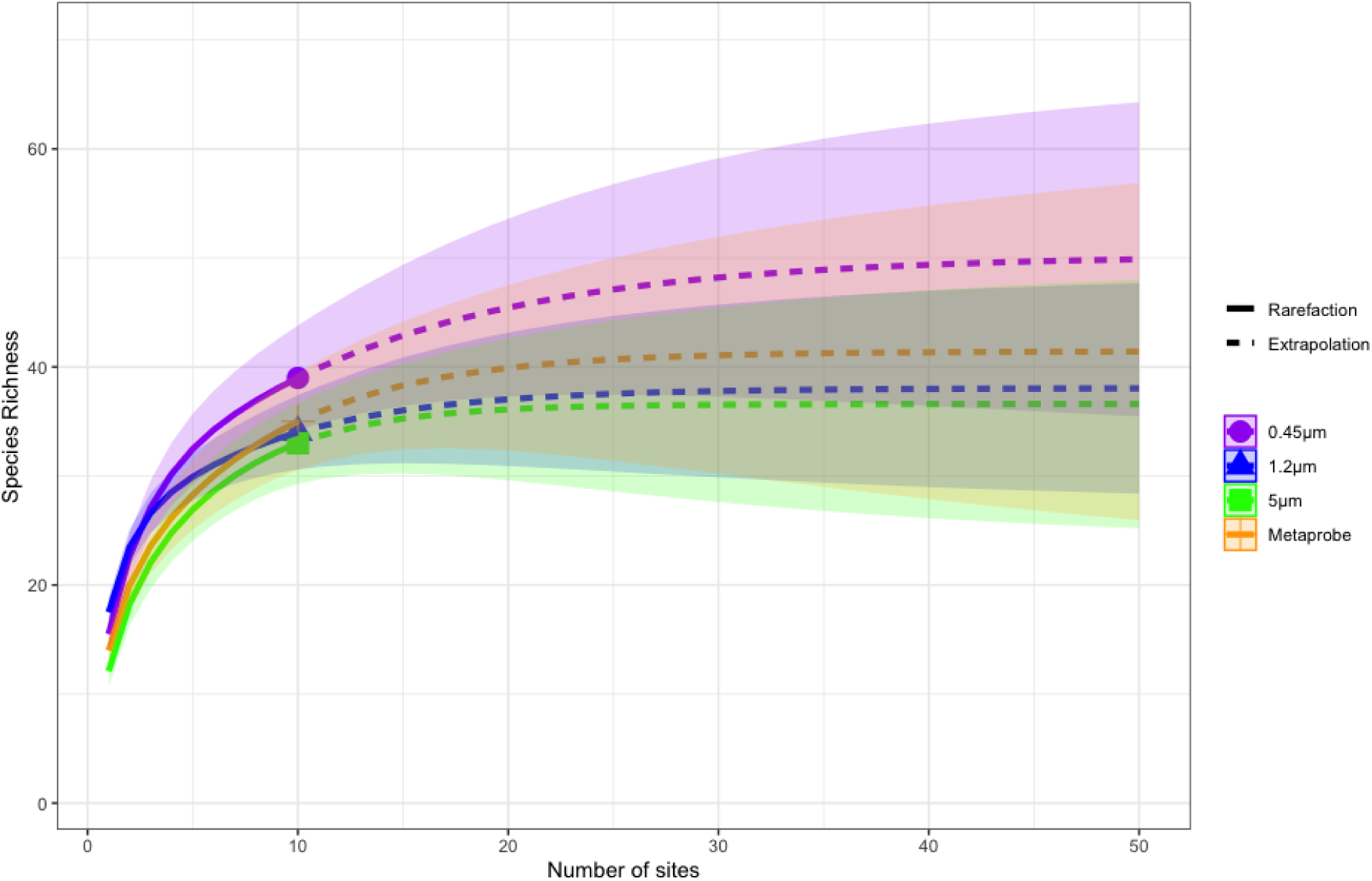
Rarefaction and extrapolation curves of species richness detected by capturing methods according to the number of sampled sites. Solid “Rarefaction” lines represent the richness detected and dotted “Extrapolation” lines represent the estimated sampling effort required based on sampling sites to reach a sampling plateau.

When analysing species richness obtained from each sampling method and between habitat types, the highest overall species richness obtained was from the upper zone using the 0.45 μm and 1.2 μm filters (Fig. 5). For each of the other zones (central and lower), the 0.45 μm and 1.2 μm filters again, recovered the highest overall species richness compared to the other methods (5 μm and metaprobe). Across the central and lower zones, the 5 μm detected the lowest overall richness. The metaprobe detected the lowest overall richness in the upper zone while detecting slightly more in the central and lower zones compared to the 5 μm. The overlaid jitter points show the variability in species detections across replicates for each sampled month. This variability is evidenced by the dispersion of points within the same colour group, indicating differences in species composition among replicates. The lack of clustering of points of the same colour suggests potential environmental heterogeneity, methodological variability, or stochasticity influencing detection rates.

**Figure 5.**
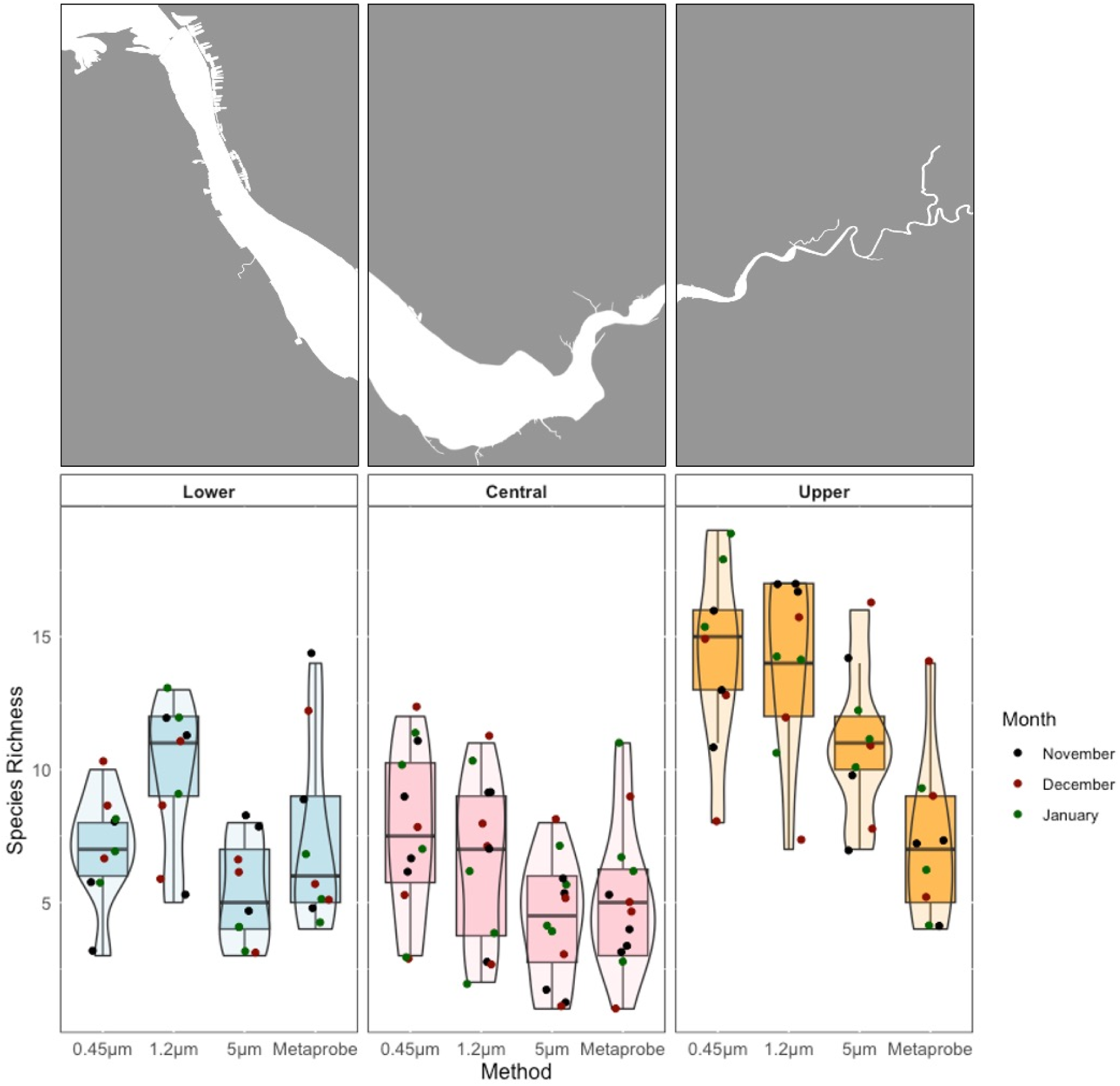
Grouped boxplot and violin plot of overall species richness for each replicate recovered across November 2022, December 2022 and January 2023 by each sampling method within a specific zone. Jittered points overlaid showing the monthly variation in species richness detection.

The GLMM was applied to our data following goodness of fit assessment using Akaike Information Criterion (AIC: 600.9) and Bayesian Information Criterion (BIC: 637.1). From the results, the 0.45 µm filtering method performs best in the upper zone and species richness detection significantly decreases in the central and lower zone (central: estimate = -0.6227, *P* = 0.0001; lower: estimate = -0.6937, *P* = 0.0001). Additional significant decreases in species richness were observed when comparing the 5 µm method in the central zone (estimate = -0.5706, *P* = 0.0009), the metaprobe in the upper zone (estimate = -0.6776, *P* = 0.000009), and the metaprobe in the central zone (estimate = -0.3947, *P* = 0.0162). Minor, non-significant increases in species richness were found using the 1.2 µm method in the lower zone (estimate = 0.3184, *P* = 0.0523) and the metaprobe in the central zone (estimate = 0.0458, *P* = 0.7930). Overall, the greatest decreases in species richness were observed with the 5 µm and metaprobe methods, while the 0.45 µm and 1.2 µm methods showed the least decreases, with no statistical difference between these two methods (Table S7).

Pearson’s correlation between volume filtered and richness found a significant correlation (Fig. S4), showing that an increase in water filtered tends to yield a higher species richness (*R* = 0.73, *P <* 0.001). The result from the SRS scaling (Fig. S5), at the normalised volume (*Cmin* = 180 ml), brings each of the three syringe capture methods closer in line with one another in terms of species richness recovered across each of the three zones, however, the same patterns are still present with 0.45 µm and the 1.2 µm showing overall consistent higher levels of richness.

The results from the analyses of species detection probabilities (Fig. 6A) indicated a higher detection probability when using the 0.45 μm for the European sprat and common bream, with the 1.2 μm method having a slightly higher detection probability for the European eel. The 0.45 μm method has a higher detection probability for all three representative species than the 5 μm and the metaprobe. The 1.2 μm shows a slightly lower detection probability than the metaprobe for the European sprat. Overall the 0.45 μm shows a more consistent and higher detection probability across the four methods tested and the three representative species. The species-specific cumulative detection probability (Fig. 6B) shows the estimated species-specific detection probabilities based on increasing sampling effort, up to a detection probability of 0.95 (dashed line). The results indicate that ∼3-5 replicates are required to reliably detect the three representative species using the 0.45 μm method, with the 1.2 μm performing similarly for two of representative species but not for the European sprat (Fig. 6B). The number of replicates required for the metaprobe was similar to the 0.45 and 1.2 μm methods for the common bream and European sprat but was markedly higher for European eel (as was the case for the 5 μm method; Fig. 6B).

**Figure 6.**
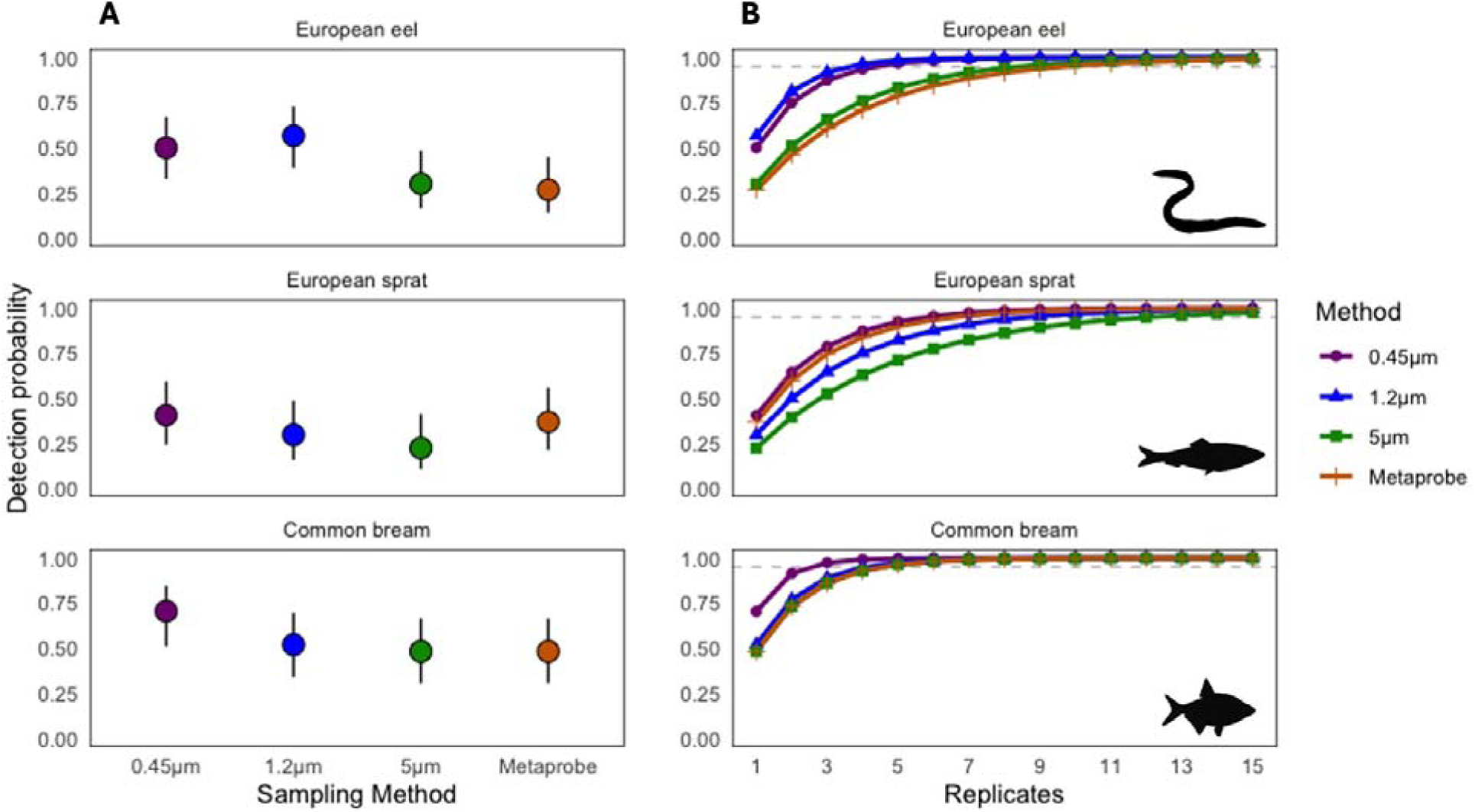
Estimated detection probabilities of each capturing method for each of the three focal species (top: European eel, *Anguilla anguilla*; middle: European sprat, *Sprattus sprattus*; bottom: common bream, *Abramis brama*) with 95% confidence intervals (A panels). Species-specific cumulative detection probability with an increasing number of sample replicates; the horizontal dashed line shows a probability of 0.95 of species detection based on the number of replicates collected (B panels).

## Discussion

The primary aim of this study was to evaluate the performance of four eDNA capture methods along a ∼60 km stretch of the Mersey estuary, a dynamic and highly heterogeneous habitat. The Mersey estuary presents unique challenges due to its macro/hyper-tidal nature and large spatio-temporal variations in water chemistry, temperature, hydrodynamics, and turbidity (Blott et al., 2006). By comparing three manual methods and one passive eDNA capturing method, we assessed their effectiveness in detecting fish diversity and discuss below their applicability for routine biomonitoring in estuarine systems. Overall, we highlight the key findings regarding capture method performance, zone-specific considerations of the study system, sensitive species detection, and practical considerations for future eDNA monitoring campaigns.

### Performance of the eDNA capture methods

The 0.45 μm capture method detected 39 species, the 1.2 μm detected 34 species, the 5 μm detected 33 species and the metaprobe detected 35 species across the three months (Fig. 3B), suggesting all methods as viable eDNA capture options. Water samples at the upper zone (freshwater) sites U1-U3 (Fig. 1) during the months sampled appeared less turbid than the seaward zones, and each sample replicate had less visible sediment present (personal observation), potentially explaining the lower filter clogging rate observed during the filtration process in this zone (Wittwer et al., 2018), and the larger volume of water filtered at each of these sites compared to the sites in the other zones (central and lower; Tables S4-S6). Overall, there was a higher species richness found in the upper zone (22) than in the central (13) and lower zones (14) and a consistently higher richness obtained by the 0.45 μm method, coinciding with previous studies suggesting a greater filtered volume will likely yield a greater richness (Lopes et al., 2017; Tsuji et al., 2019; Bessey et al., 2020; Schabacker et al., 2020). This is further substantiated by a clear correlation between volume filtered and species richness (Fig. S4), and the statistical differences observed via the pairwise PERMANOVA test (Table S3). The SRS normalisation of richness based on a minimum volume (*Cmin* = 180 ml) (Fig. S5) reveals that richness recovered by each method became more consistent across methods when normalised to an equal volume. However, disparities in species richness between the zonal salinity gradients remain, particularly in the upper zone where a higher richness persists (Fig. S5). As discussed in earlier studies (Stoeckle et al., 2017; Barnes et al., 2021; Altermatt et al., 2023), it should be considered that volume may not be the sole determinant in recovering higher species richness estimates. Ecological factors, such as habitat structure, salinity gradients, and the connectivity between environmental zones, can significantly influence species richness (Leibold et al., 2004; Lin et al., 2024) and other methodological considerations, including sampling effort, the taxonomic resolution of primers and reference databases, also play key roles in determining species recovery (Ruppert et al., 2019). Therefore, the selection of an optimal method may require consideration of not only the highest richness recovered but also consistency in performance.

The GLMM results presented here give an insight into the performance and consistency of the four eDNA sampling methods. The results indicate significant decreases (*P* = 0.0009) when using the 5 μm method in the central zone compared to the 0.45 μm, which may be a contributing factor to the richness obtained, as smaller DNA fragments may not be captured by the larger pore size (Turner et al., 2014; Barnes et al., 2021; Jo, Takao & Minamoto, 2022) and more significant decreases in the upper (*P* = 0.0162) and central (*P* = 0.000008) zones compared to the 0.45 μm. We do, however, see increases (but non-significant) in species richness when using the 1.2 μm method in the lower zone over the 0.45 μm method, and when using the metaprobe in the lower zone (Table S6). The key result is that the most consistent performing filter types were the 0.45 µm and the 1.2 µm, with the two showing the highest amount of richness across all zones compared to the other filter types. Interpretation of this data should be underpinned by the fact that the richness recovered is based on eDNA presence, and not the presence of a physical organism (Bohmann et al., 2014), therefore, due to the samples being collected at high tide only, and this study system being characterised by strong tidal movements (Bowden, 1963), this could have resulted in high levels of eDNA transport across the sampled zones through hydrological behaviour (Deiner & Altermatt, 2014; Ruppert et al., 2019; Laporte et al., 2022). As a result, this could lead to false positives of species detections in certain zones and sites (e.g., freshwater species detected in marine locations and marine species detected in freshwater locations; Burian et al., 2021).

Previous studies have shown how sampling effort (through duration or replicate numbers) can improve species detections, particularly in aquatic environments (Willoughby et al., 2016; Millard-Martin et al., 2024). This was evident here when looking at the sampling effort of three replicates per method used at each of the 10 sites, and it is suggested that there is scope for increased species detections for each method (Fig. 5). The rarefaction curves for the 1.2 μm, 5 μm, and metaprobe capture methods indicate that species detections could improve with increased sampling effort, potentially reaching around ∼40 species. However, the curves reach a plateau at approximately 30 sites, with only minor increases observed beyond this point. The 0.45 µm method suggests only a slightly higher extrapolated richness than the observed (44) with effort increased to ∼40-50 sites. These additional species detections for all four methods would require a drastic increase in the sampling effort carried out (10 sites), which could be considered counterintuitive. This highlights that while increased sampling effort spatially with additional sites may enhance species richness estimates, the gains may not justify the substantial increase in effort required. Furthermore, as was present in this study, access restrictions or challenging terrain may dictate the number of sample sites, therefore, a greater sampling effort here may be more applicable through an increase in the number of replicates taken for all methods (Furlan et al., 2016; Lawson Handley et al., 2019), and time deployed for the metaprobe (e.g., five replicates instead of three; 30 minutes instead of 5).

Accuracy and repeatability are critical factors in eDNA studies, especially if a particular species is the aim of detection within a study system (Mauvisseau et al., 2019; Mathieu et al., 2020). Species detection probabilities were assessed using the four methods and three replicates, to provide an insight into how consistent each method was at detecting a particular species (Fig. 6). We used three representative species, which gave us a balance of species detected across all zones (*A. anguilla* European eel), detected in the lower zone (*O. eperlanus* European sprat) and detected in the upper zone (*A. brama* common bream). When looking at the detection probabilities (Fig. 6A), the 0.45 μm method had a more consistent detection rate (45-75%) for all species examined, whereas the other three methods had more varied detection rates (1.2 μm filters: 25-60%; 5 μm: 20-50%; metaprobe: 20-50%), indicating a lower level of consistency compared to the 0.45 μm method. Cumulative detection probability (Fig. 6B) suggests that if the number of replicates for the 0.45 μm filter was increased to five, there would be a higher likelihood of achieving a 0.95 probability of detecting all three representative species. This result reinforces the consistent reports of increased effort yielding more data (Erickson et al., 2019; Yates et al., 2023). When considered alongside the findings in Fig. 4, which also indicate the potential need for increased sampling effort, for this study system and similar systems worldwide, we recommend an increase in the sampling effort used here (10 sites and five replicates at each).

The application of eDNA monitoring approaches for low population/rare species detections has been widely documented (Jerde et al., 2011; Beng & Corlett, 2020; Duarte et al., 2023). Within the data, four UK Biodiversity Action Plan (BAP) priority fish species are present: the European eel, brown trout, Atlantic salmon and the European smelt (Joint Nature Conservation Committee, 2007). Of these species, it is estimated that the Atlantic salmon population numbers in the Mersey are extremely low (Ikediashi et al., 2012), and the population numbers of the European smelt are entirely unknown but also assumed to be low as there are currently no catch records to generate an estimation. Out of the four eDNA capturing methods, only the 0.45 μm successfully detected all four species (Table S2). The European eel and brown trout were detected by all methods, whereas the Atlantic salmon was detected by the 0.45 μm, 1.2 μm and metaprobe but not detected at all by the 5 μm method. The European smelt is exclusively detected by the 0.45 μm method. Detecting all four species present in the data may indicate the 0.45 μm method as being a more effective approach, especially if the detection of low-population/rare species is a key aspect of the study design (Sepulveda et al., 2019).

### Usability and practicality of each capture method

As indicated in earlier eDNA research, the efficacy of an eDNA protocol should be thoroughly assessed prior to deployment (Piggot, 2016; Diaz et al., 2020; Kumar et al., 2020), particularly in complex estuarine systems, where challenges such as fluctuating salinity, tidal variability, and nutrient dynamics may influence species interactions in ways not typically encountered in other ecosystem types (Matheson et al., 2010). Here, a key consideration was consistency in the ability of the syringe methods to process water across all sampled zones without becoming clogged prematurely, with the aim of filtering as much water as possible per method to provide robust results (Turner et al., 2014; Thomas et al., 2018). Although the filter method pore size increased to 1.2 μm and 5 μm, the increase in pore size did not allow for larger volumes of water to be filtered, with more water consistently filtered using the 0.45 μm method (Tables S4-S6). One potential explanatory reason for this was a noticeable difference between the filter designs. The Sterivex 0.45 μm filters have a cylindrical form, with excess space for around ∼3 ml of water to be held within the filter cartridge, whereas the Whatman 1.2 μm and the KX Nylon 5 μm filters used here are disc form filters, with no excess space in the filter cartridge. The extra space within the Sterivex filters may allow room for some suspended sediments to remain suspended, as opposed to the disc form filters which applied all suspended sediments directly onto the filter membrane immediately, thus potentially explaining the filter clogging at a faster rate. In addition, it was found that laboratory processing of the 0.45 μm filter cartridges was more simplistic than the other filters, due to the design of the product (the presence of an easily removable cap, not present in the 1.2 μm and the 5 μm filters used here). The 1.2 μm and 5 μm filter cartridges had to be cut open with excessive force using wire cutters, which added effort and time to the process, thus making the 0.45 μm easier to use. Moreover, DNA extraction methods which are performed within the syringe cartridge without removing the filter membrane (Miya et al, 2016; Bowen et al, 2024), may be better suited to the 0.45 μm method due to the extra space present, providing potentially more flexibility in its use. The 1.2 μm and 5 μm filters used here do not have any excess space and, therefore, this option would need assessing to determine if the physical design of these filter types would allow for this type of extraction method. Therefore, a key consideration to be taken into account before sampling in similar systems, in addition to pore size, should be filter design.

When using the metaprobe, any issues experienced with filter clogging are eliminated, as there is no manual or mechanical filtration through a filter cartridge and membrane (Maiello et al., 2022, 2024). As discussed in earlier research, moving towards passive eDNA collection methods can reduce field time, manual labour and remove a need for specialised equipment such as filters and pumps (Mariani et al., 2019; Bessey et al., 2021), which leads to key benefits of this method, making it a potentially more accessible approach (Bessey et al., 2022). Passive samplers have been increasingly utilised over recent years, yielding positive results (Chen et al., 2022; van der Hayde et al., 2023; Cai et al., 2024), which is further substantiated here, with this being the first known dataset highlighting the metaprobe as an effective passive eDNA sampling method within estuarine and freshwater river systems. We deployed the metaprobe for a short period of time (five minutes) to coincide with the manual filtration time frames, and the metaprobe detected a high number of species in comparison to the other methods (35 total). The short deployment time and high number of species detected again, support the claims of passive sampling becoming a highly viable eDNA sampling approach (Jeunen et al., 2024).

A drawback of using the metaprobe over the more commonly used syringe/filter-based methods is that the volume of water that has passed through the metaprobe is challenging to quantify, and would need further investigation and testing. Therefore, any potential control over improving species detection rates through avenues such as filtering larger volumes of water, more simply implemented with a syringe-based method (Hunter et al., 2019), cannot be performed as easily. Options to increase species detections could come from an increase in sample replicates within the metaprobe and/or time deployed at sample sites. Furthermore, a potential optimisation approach which may apply to this method would be to assess the water flow rate within a study system before the metaprobe deployment, targeting periods of higher flow in which data via tide gauges can be freely accessed, and can be used to predict tidal behaviour (Hibbert et al., 2015). Given that this method relies on water passively moving through the metaprobe, it would be worthwhile to test if there is a correlation between higher flow rates and species richness. In addition, this method may present some barriers to use, as it does require 3D printing of the probes, which may not be an easily accessible option for all end users. Furthermore, the gauze material once it has been used to capture eDNA requires a storage solution, typically ethanol (Maiello et al., 2022, 2024; Neave, Mariani & Meek, 2023). Currently, there is no data available comparing storage solutions for alternative approaches to ethanol, which again may cause issues in the accessibility of ethanol for purchase, transport and use in the field. This, then needs to be carefully planned and considered before use.

## Conclusions

This study demonstrates that eDNA metabarcoding can be applied successfully for biomonitoring in heterogeneous estuarine systems. The four methods tested obtained a high number of species detections, ranging from 39 species detected (0.45 µm Sterivex filter) to 33 (5 µm KX Nylon filter). The 0.45 µm filter was considered the optimal choice, as it achieved the most consistent results across the ten sampling sites throughout the estuary by yielding the highest richness estimates, detecting all UK BAP species present, and producing the least variability in species detection probabilities. However, all four methods were considered viable eDNA capture methods, and if budget is a constraint, then the metaprobe would be an effective, easy-to-use alternative option. Lastly, in highly dynamic estuarine systems, such as the Mersey, tidal patterns directly influence the transport of eDNA particles, which, in turn, should be considered when interpreting the results. This emphasises the need for integrative approaches, such as including hydrodynamic modelling and particle tracking methods to simulate hydrological patterns and the subsequent dispersal of eDNA (Andruszkiewicz et al., 2019; Pont, 2024). This would allow for more accurate interpretation of eDNA results, better identification of actual species presence at a particular location, and contribute towards building more effective biomonitoring strategies.

## Data Availability Statement

The data that support the findings of this study are openly available in - JmJackman27/Mersey_eDNA_data – at https://github.com/JmJackman27/Mersey_eDNA_data.git.

