## Supplementary material for "Performance of eDNA capture methods for monitoring fish biodiversity in a hyper-tidal estuary"

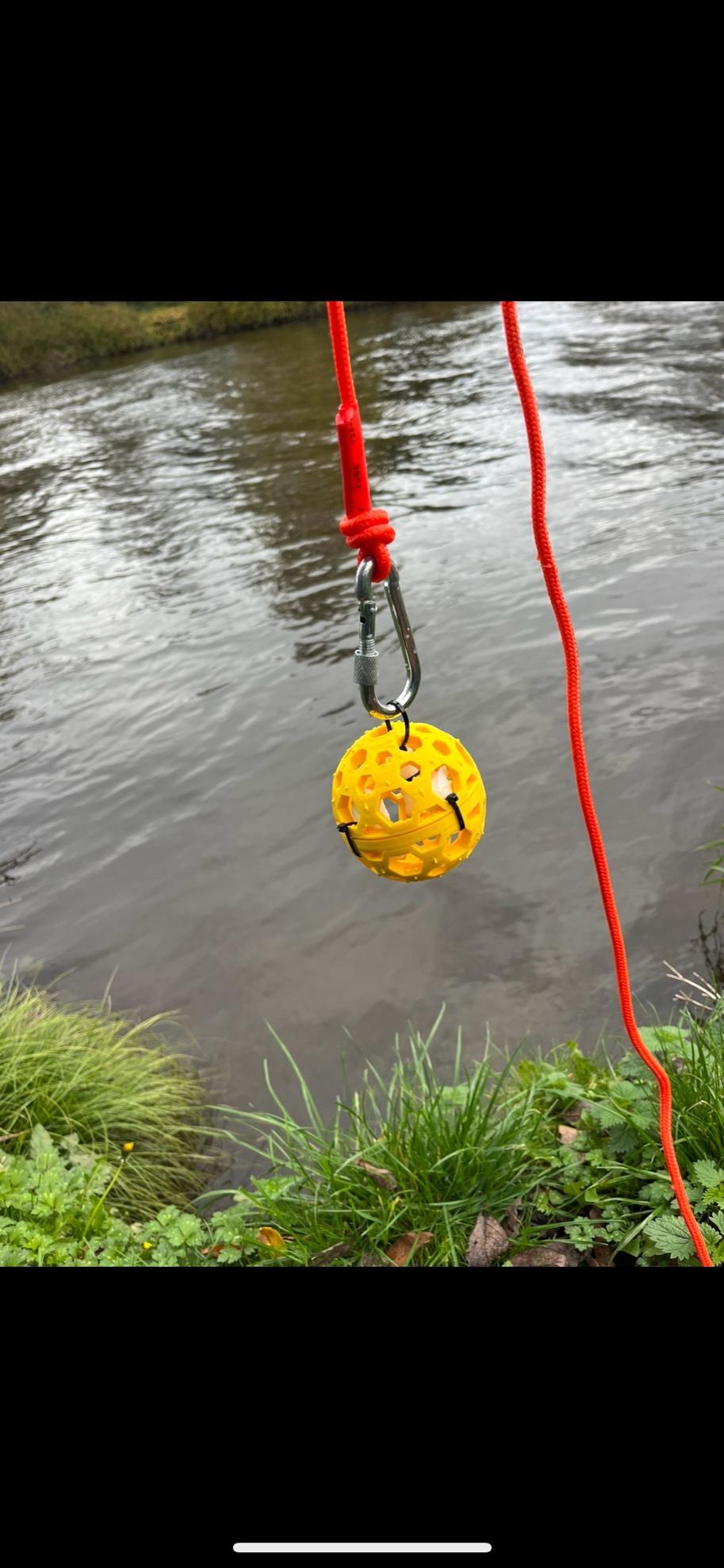

Figure S1. The metaprobe, attached to a 20 m rope with a

Carabiner to be cast out at each sampling site.

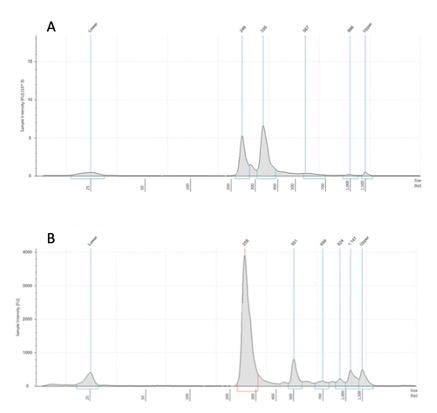

Figure S2. Comparison of amplification of target fragments for two fish-specific primer sets. (A) is the MiFish primer set and (B) is the Tele02 primer set using the Agilent 2200 tapestation with D1000 High sensitivity screentape.

**Table S1.** Filtering steps removing all MOTUs/reads originating from sequencing errors or contamination and respective number of reads retrieved at each stage.

| **Filtering Steps** | **Total** |
| --- | --- |
| **Total Reads** | **49,529,742** |
| After removing reads from the blanks and positive controls | 47,997,886 |
| After removing all non-fish reads | 6,131,765 |

**Table S2**. Overall species list of all detections across all filtering methods, sample sites and dates.

| **Order** | **Family** | **Species** | **Common name** | **Detection method** |
| --- | --- | --- | --- | --- |
| Anguilliformes | Anguillidae | *Anguilla anguilla* | European eel | 0.45µm, 1.2µm, 5µm, Metaprobe |
| Blenniiformes | Blenniidae | *Lipophrys pholis* | Shanny | 0.45µm, 1.2µm, 5µm |
| Clupeiformes | Clupeidae | *Clupea harengus* | Atlantic herring | 0.45µm, 1.2µm, 5µm, Metaprobe |
| Clupeiformes | Clupeidae | *Sprattus sprattus* | European sprat | 0.45µm, 1.2µm, 5µm, Metaprobe |
| Clupeiformes | Alosidae | *Sardina pilchardus* | European pilchard | 1.2µm, 5µm, Metaprobe |
| Cypriniformes | Cyprinidae | *Rutilus rutilus* | Common roach | 0.45µm, 1.2µm, 5µm, Metaprobe |
| Cypriniformes | Cyprinidae | *Abramis brama* | Common bream | 0.45µm, 1.2µm, 5µm, Metaprobe |
| Cypriniformes | Cyprinidae | *Phoxinus phoxinus* | Common minnow | 0.45µm, 1.2µm, 5µm, Metaprobe |
| Cypriniformes | Cyprinidae | *Gobio gobio* | Gudgeon | 0.45µm, 1.2µm, 5µm, Metaprobe |
| Cypriniformes | Cyprinidae | *Squalius cephalus* | Chub | 0.45µm, 1.2µm, 5µm, Metaprobe |
| Cypriniformes | Cyprinidae | *Leuciscus leuciscus* | Dace | 0.45µm, 1.2µm, 5µm, Metaprobe |
| Cypriniformes | Nemacheilidae | *Barbatula barbatula* | Stone loach | 0.45µm, 1.2µm, 5µm, Metaprobe |
| Cypriniformes | Cyprinidae | *Carassius carassius* | Crucian carp | 0.45µm |
| Cypriniformes | Cyprinidae | *Blicca bjoerkna* | White bream | 0.45µm, 1.2µm, 5µm, Metaprobe |
| Cypriniformes | Cyprinidae | *Cyprinus carpio* | Common carp | 0.45µm |
| Cypriniformes | Cyprinidae | *Scardinius erythrophthalmus* | Common rudd | 0.45µm, 1.2µm |
| Esociformes | Esocidae | *Esox lucius* | Northern pike | 0.45µm, 1.2µm, 5µm, Metaprobe |
| Gadiformes | Gadidae | *Merlangius merlangus* | Whiting | 0.45µm, 1.2µm, 5µm, Metaprobe |
| Gadiformes | Lotidae | *Ciliata mustela* | Fivebeard rockling | 0.45µm, 1.2µm, 5µm, Metaprobe |
| Gadiformes | Gadidae | *Gadus morhua* | Atlantic cod | 0.45µm, 1.2µm, 5µm |
| Gadiformes | Gadidae | *Trisopterus minutus* | Poor cod | 5µm, Metaprobe |
| Gasterosteiformes | Gasterosteidae | *Gasterosteus aculeatus* | Three-spined stickleback | 0.45µm, 1.2µm, 5µm, Metaprobe |
| Gobiiformes | Gobiidae | *Pomatoschistus microps* | Common goby | 0.45µm, 1.2µm, 5µm, Metaprobe |
| Gobiiformes | Gobiidae | *Pomatoschistus minutus* | Sand goby | 0.45µm, 1.2µm, 5µm, Metaprobe |
| Moroniformes | Moronidae | *Dicentrarchus labrax* | European seabass | 0.45µm, 1.2µm, 5µm, Metaprobe |
| Mugiliformes | Mugilidae | *Chelon labrosus* | Thicklip grey mullet | 0.45µm, Metaprobe |
| Osmeriformes | Osmeridae | *Osmerus eperlanus* | European smelt | 0.45µm |
| Perciformes | Percidae | *Perca fluviatilis* | European perch | 0.45µm, 1.2µm, 5µm, Metaprobe |
| Perciformes | Percidae | *Gymnocephalus cernua* | Eurasian ruffe | 0.45µm, 1.2µm, 5µm, Metaprobe |
| Perciformes | Sparidae | *Sparus aurata* | Gilt-head bream | 0.45µm, Metaprobe |
| Pleuronectiformes | Pleuronectidae | *Platichthys flesus* | European flounder | 0.45µm, 1.2µm, 5µm, Metaprobe |
| Pleuronectiformes | Soleidae | *Solea solea* | Common sole | 0.45µm, 1.2µm, 5µm, Metaprobe |
| Pleuronectiformes | Pleuronectidae | *Limanda limanda* | Common dab | 0.45µm, 5µm, Metaprobe |
| Pleuronectiformes | Pleuronectidae | *Pleuronectes platessa* | European plaice | 0.45µm |
| Pleuronectiformes | Soleidae | *Buglossidium luteum* | Yellow sole | 5µm, Metaprobe |
| Salmoniformes | Salmonidae | *Salmo salar* | Atlantic salmon | 0.45µm, Metaprobe |
| Salmoniformes | Salmonidae | *Salmo trutta* | Brown trout | 0.45µm, 1.2µm, 5µm, Metaprobe |
| Salmoniformes | Salmonidae | *Thymallus thymallus* | European grayling | 0.45µm, 1.2µm, 5µm, Metaprobe |
| Salmoniformes | Salmonidae | *Oncorhynchus gorbuscha* | Pink salmon | 0.45µm |
| Scombriformes | Scombrini | *Scomber scombrus* | Atlantic mackerel | 0.45µm, 1.2µm, 5µm, Metaprobe |
| Scorpaeniformes | Cottidae | *Cottus gobio* | European bullhead | 0.45µm, 1.2µm, 5µm, Metaprobe |
| Scorpaeniformes | Gasterosteidae | *Pungitius pungitius* | Ninespine stickleback | 1.2µm |
| Sygnathiformes | Syngathidae | *Syngnathus rostellatus* | Lesser pipefish | 1.2µm, 5µm, Metaprobe |
| Trachiniformes | Ammodytidae | *Ammodytes marinus* | Raitts’s sand eel | 0.45µm, 1.2µm, Metaprobe |

**Table S3. PERMANOVA pairwise comparisons of richness between method**

| **Comparison** | **Df** | **Sum Of Sqs** | **R^2^** | **F value** | **P value** | **Padjusted** | **PSig** |
| --- | --- | --- | --- | --- | --- | --- | --- |
| 0.45µm vs 1.2µm | 1 | 0.0057765 | 0.00184702 | 0.10732561 | 0.869 | 1 |  |
| 0.45µm vs 5µm | 1 | 0.37233268 | 0.08985982 | 5.72644719 | 0.007 | 0.042 | * |
| 0.45µm vs Metaprobe | 1 | 0.38629367 | 0.11240431 | 7.34506733 | 0.004 | 0.024 | * |
| 1.2µm vs 5µm | 1 | 0.41421059 | 0.09381099 | 6.0043077 | 0.005 | 0.03 | * |
| 1.2µm vs Metaprobe | 1 | 0.44632917 | 0.11976549 | 7.89153194 | 0.004 | 0.024 | * |
| 5µm vs Metaprobe | 1 | 0.01903599 | 0.00482063 | 0.28095075 | 0.767 | 1 |  |

**Table S4: Volumes of water filtered per sample replicate for November 2022**

| **Volume filtered in millilitres** | | | | |
| --- | --- | --- | --- | --- |
| Site | Filter | Rep_1 | Rep_2 | Rep_3 |
| U1 | 0.45μm | 600 | 540 | 580 |
| U1 | 1.2μm | 540 | 550 | 530 |
| U1 | 5μm | 320 | 360 | 320 |
| U2 | 0.45μm | 500 | 560 | 480 |
| U2 | 1.2μm | 400 | 400 | 420 |
| U2 | 5μm | 340 | 310 | 300 |
| U3 | 0.45μm | 420 | 450 | 440 |
| U3 | 1.2μm | 400 | 420 | 380 |
| U3 | 5μm | 300 | 310 | 300 |
| C1 | 0.45μm | 480 | 440 | 440 |
| C1 | 1.2μm | 390 | 400 | 360 |
| C1 | 5μm | 320 | 300 | 320 |
| C2 | 0.45μm | 420 | 380 | 310 |
| C2 | 1.2μm | 290 | 300 | 290 |
| C2 | 5μm | 200 | 220 | 240 |
| C3 | 0.45μm | 340 | 380 | 320 |
| C3 | 1.2μm | 300 | 360 | 320 |
| C3 | 5μm | 220 | 200 | 190 |
| C4 | 0.45μm | 400 | 380 | 410 |
| C4 | 1.2μm | 380 | 350 | 380 |
| C4 | 5μm | 280 | 200 | 260 |
| L1 | 0.45μm | 500 | 450 | 420 |
| L1 | 1.2μm | 400 | 450 | 390 |
| L1 | 5μm | 340 | 280 | 300 |
| L2 | 0.45μm | 420 | 440 | 400 |
| L2 | 1.2μm | 380 | 340 | 400 |
| L2 | 5μm | 320 | 300 | 290 |
| L3 | 0.45μm | 460 | 410 | 380 |
| L3 | 1.2μm | 360 | 440 | 400 |
| L3 | 5μm | 250 | 220 | 280 |

**Table S5: Volumes of water filtered per sample replicate for December 2022**

| **Volume filtered in millilitres** | | | | |
| --- | --- | --- | --- | --- |
| Site | Filter | Rep_1 | Rep_2 | Rep_3 |
| U1 | 0.45μm | 650 | 560 | 550 |
| U1 | 1.2μm | 500 | 460 | 480 |
| U1 | 5μm | 400 | 420 | 400 |
| U2 | 0.45μm | 520 | 500 | 460 |
| U2 | 1.2μm | 400 | 450 | 440 |
| U2 | 5μm | 380 | 400 | 360 |
| U3 | 0.45μm | 520 | 460 | 550 |
| U3 | 1.2μm | 460 | 400 | 450 |
| U3 | 5μm | 380 | 330 | 340 |
| C1 | 0.45μm | 420 | 360 | 380 |
| C1 | 1.2μm | 380 | 380 | 350 |
| C1 | 5μm | 300 | 260 | 280 |
| C2 | 0.45μm | 350 | 380 | 380 |
| C2 | 1.2μm | 360 | 310 | 320 |
| C2 | 5μm | 280 | 220 | 230 |
| C3 | 0.45μm | 300 | 220 | 260 |
| C3 | 1.2μm | 260 | 300 | 250 |
| C3 | 5μm | 200 | 210 | 160 |
| C4 | 0.45μm | 360 | 330 | 370 |
| C4 | 1.2μm | 300 | 320 | 350 |
| C4 | 5μm | 260 | 280 | 210 |
| L1 | 0.45μm | 330 | 340 | 340 |
| L1 | 1.2μm | 300 | 280 | 300 |
| L1 | 5μm | 220 | 250 | 260 |
| L2 | 0.45μm | 300 | 310 | 270 |
| L2 | 1.2μm | 340 | 260 | 300 |
| L2 | 5μm | 250 | 220 | 240 |
| L3 | 0.45μm | 380 | 350 | 360 |
| L3 | 1.2μm | 330 | 300 | 310 |
| L3 | 5μm | 260 | 220 | 240 |

**Table S6: Volumes of water filtered per sample replicate for January 2023**

| **Volume filtered in millilitres** | | | | |
| --- | --- | --- | --- | --- |
| Site | Filter | Rep_1 | Rep_2 | Rep_3 |
| U1 | 0.45μm | 520 | 510 | 480 |
| U1 | 1.2μm | 520 | 500 | 450 |
| U1 | 5μm | 400 | 380 | 450 |
| U2 | 0.45μm | 550 | 520 | 460 |
| U2 | 1.2μm | 450 | 410 | 370 |
| U2 | 5μm | 380 | 360 | 340 |
| U3 | 0.45μm | 490 | 500 | 450 |
| U3 | 1.2μm | 460 | 440 | 400 |
| U3 | 5μm | 300 | 380 | 320 |
| C1 | 0.45μm | 460 | 390 | 400 |
| C1 | 1.2μm | 410 | 400 | 310 |
| C1 | 5μm | 260 | 220 | 300 |
| C2 | 0.45μm | 300 | 290 | 300 |
| C2 | 1.2μm | 270 | 220 | 250 |
| C2 | 5μm | 210 | 200 | 190 |
| C3 | 0.45μm | 330 | 350 | 300 |
| C3 | 1.2μm | 270 | 310 | 280 |
| C3 | 5μm | 200 | 180 | 180 |
| C4 | 0.45μm | 360 | 340 | 380 |
| C4 | 1.2μm | 300 | 260 | 240 |
| C4 | 5μm | 180 | 180 | 200 |
| L1 | 0.45μm | 460 | 410 | 440 |
| L1 | 1.2μm | 380 | 390 | 400 |
| L1 | 5μm | 300 | 290 | 240 |
| L2 | 0.45μm | 400 | 370 | 380 |
| L2 | 1.2μm | 330 | 360 | 320 |
| L2 | 5μm | 220 | 210 | 180 |
| L3 | 0.45μm | 360 | 400 | 350 |
| L3 | 1.2μm | 240 | 300 | 270 |
| L3 | 5μm | 220 | 200 | 180 |

**Table S7.** GLMM results showing richness decreases and increases between different eDNA capture methods across different zones within the Mersey estuary.
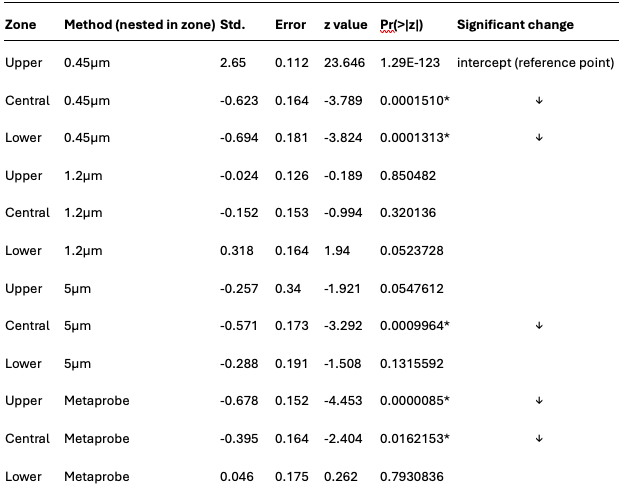

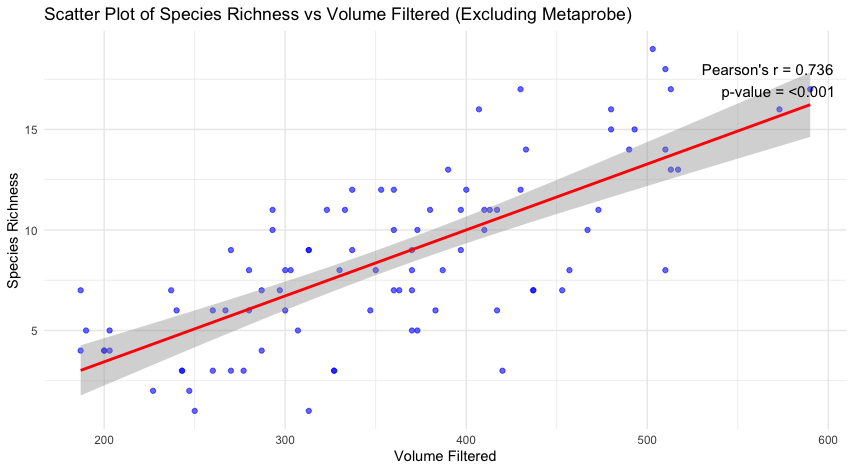

Figure S4. The correlation between species richness and the volume of water filtered indicates a strong relationship between a higher volume of water filtered yielding a higher level of species richness.

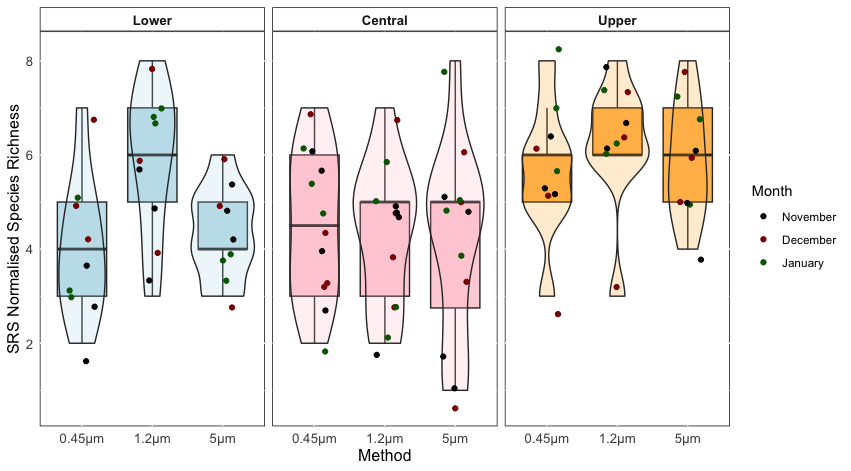

Figure S5. Scaling ranked subsampling of species richness based on *Cmin* =180ml for the 0.45µm, 1.2µm and 5µm eDNA capture methods.
